# The endoplasmic reticulum connects to the nucleus by constricted junctions that mature after open mitosis in mammalian cells

**DOI:** 10.1101/2023.01.31.526419

**Authors:** Helena Bragulat-Teixidor, Keisuke Ishihara, Gréta Martina Szücs, Shotaro Otsuka

## Abstract

The endoplasmic reticulum (ER) is physically connected to the nucleus by junctions with the outer membrane of the nuclear envelope (NE). The ER–NE junctions are essential for supplying the NE with lipids and proteins synthesized in the ER. However, little is known about the structure of these ER–NE junctions. Here, we systematically studied the ultrastructure of ER–NE junctions in cryo-fixed mammalian cells staged in anaphase, telophase, and interphase by correlating live cell imaging with three-dimensional electron microscopy. Strikingly, our results revealed that ER–NE junctions in interphase cells have a pronounced hourglass shape with a constricted neck of 7–20 nm width. This morphology is significantly distinct from that of junctions among the ER, and it emerges as early as telophase. The highly constricted ER–NE junctions are seen in several mammalian cell types, but not in budding yeast. We speculate that the unique and highly-constricted ER–NE junctions are regulated via novel mechanisms that contribute to ER-to-NE lipid and protein traffic in higher eukaryotes.

## INTRODUCTION

The endoplasmic reticulum (ER) of the eukaryotic cell is the major site of lipid and membrane protein synthesis. The ER extends from the nuclear envelope as a continuous membranous organelle, consisting of a network of tubules and sheets that are interconnected by numerous junctions (referred to as ER–ER junctions) (Nixon-Abell et al., 2016; Takakura et al., 2017). The ER makes extensive contacts with other membrane-bound organelles including mitochondria, the Golgi apparatus, and endosomes. These contact sites play essential roles in protein homeostasis, lipid transfer, metabolite shuffling, ion exchange, and signal transduction (Rossini et al., 2021). In these contact sites, ER membranes are tethered to, but not fused with the membranes of other organelles. In contrast, the ER membrane contacts the nucleus by direct fusion to the outer nuclear membrane (ONM) of the nuclear envelope (NE) (West et al., 2011). We refer to these fusion contacts as ER–NE junctions. Previous studies have shown that NE proteins can diffuse from the ER to the NE (Zuleger et al., 2011) and that NE proteins accumulate at the ER when their transport to the NE is blocked (Boni et al., 2015; Ungricht et al., 2015). Therefore, despite the presence of ribosomes at the ONM, the majority of NE proteins are expected to be synthesised on the ER and subsequently transported to the NE through ER–NE junctions.

ER–NE junctions were first visualized by electron microscopy more than 60 years ago in chemically fixed cells of rat spleen and maize rood (Watson, 1955; Whaley et al., 1960). In these cells, the ER–NE junctions had a non-constricted wide cone-shaped base that resembled the morphology of ER–ER junctions (50–100 nm). On the contrary, narrow and constricted ER–NE junctions (25-30 nm in wide) were sporadically observed in the late 1980s in high-pressure frozen plant root tip cells (Craig and Staehelin 1988; Staehelin, 1997). High-pressure freezing preserved intracellular membranes in a native state much better than chemical fixation (Dahl and Staehelin, 1989), but the finding of constricted ER–NE junctions in plant cells in the late 1980s was based on a few images of two-dimensional EM. The lack of volumetric high-resolution EM techniques at that time precluded a generalizable and systematic understanding of the three-dimensional (3D) ultrastructure of ER–NE junctions. In the early 2010s, ER–NE junctions were observed in 3D with EM tomography in high-pressure frozen yeast cells, and their morphology was reported to be similar to that of ER–ER junctions (Friedman and Voeltz, 2011). Nowadays, ER–NE junctions in all eukaryotic cells are thought to exhibit the same morphology as ER–ER junctions. Nonetheless, studies on the native ultrastructure of ER– NE junctions in high-pressure frozen mammalian cells are missing. Further, it remains unknown how ER–NE junctions are established and maintained during the cell cycle of open mitotic cells in which the NE reforms from ER membranes after having disassembled during mitosis (Ungricht & Kutay, 2017). A systematic ultrastructural analysis of ER–NE junctions in high-pressure frozen cells is required for a generalizable understanding of the 3D ultrastructure of ER–NE junctions and their biogenesis.

In this study, we combined live-cell imaging, high-pressure freezing, and quantitative 3D EM to study systematically the ultrastructure of ER–NE junctions in mammalian cells in their native state. Strikingly, we discovered that ER–NE junctions have a constricted ultrastructure that is morphologically distinct from the junctions within the ER. By correlating live-cell imaging with electron tomography, we found that the ultrastructure of ER–NE junctions is remodelled during the cell cycle, and that the membrane constriction of ER–NE junctions starts in early telophase. These findings suggest that ER–NE junctions are regulated by a novel mechanism distinct from the one remodelling junctions within the ER. The highly curved and constricted ultrastructure of ER–NE junctions likely reflects their function as sites of lipid and protein traffic between the ER and nuclear membranes, which may regulate nuclear functions.

## RESULTS

### An electron microscopy assay to study ER–NE junctions systematically throughout the cell cycle

To investigate the ultrastructure and abundance of ER–NE junctions at high temporal resolution throughout the cell cycle, we established a workflow that combines live-cell imaging with high-resolution 3D EM (Figure 1). Briefly, HeLa cells growing on carbon-coated sapphire discs were synchronized by using a double thymidine block protocol, and their progression into mitosis was monitored by time-lapse light microscopy. When most dividing cells in the field of view reached the late anaphase or the telophase stage of mitosis, we rapidly froze the discs in a high-pressure freezer. Following freeze-substitution and infiltration in plastic resin, 250 nm sections were cut, and the cells identified by time-lapse imaging were imaged by transmission EM (Figure 1A). Using this workflow, we can study the ultrastructure of cells and their organelles at precisely defined stages of the cell cycle. In this study, we focused on three time points: late anaphase (4–6 minutes after anaphase onset), early telophase (8–10 minutes after anaphase onset), and interphase (Figure 1A).

**Figure 1.**
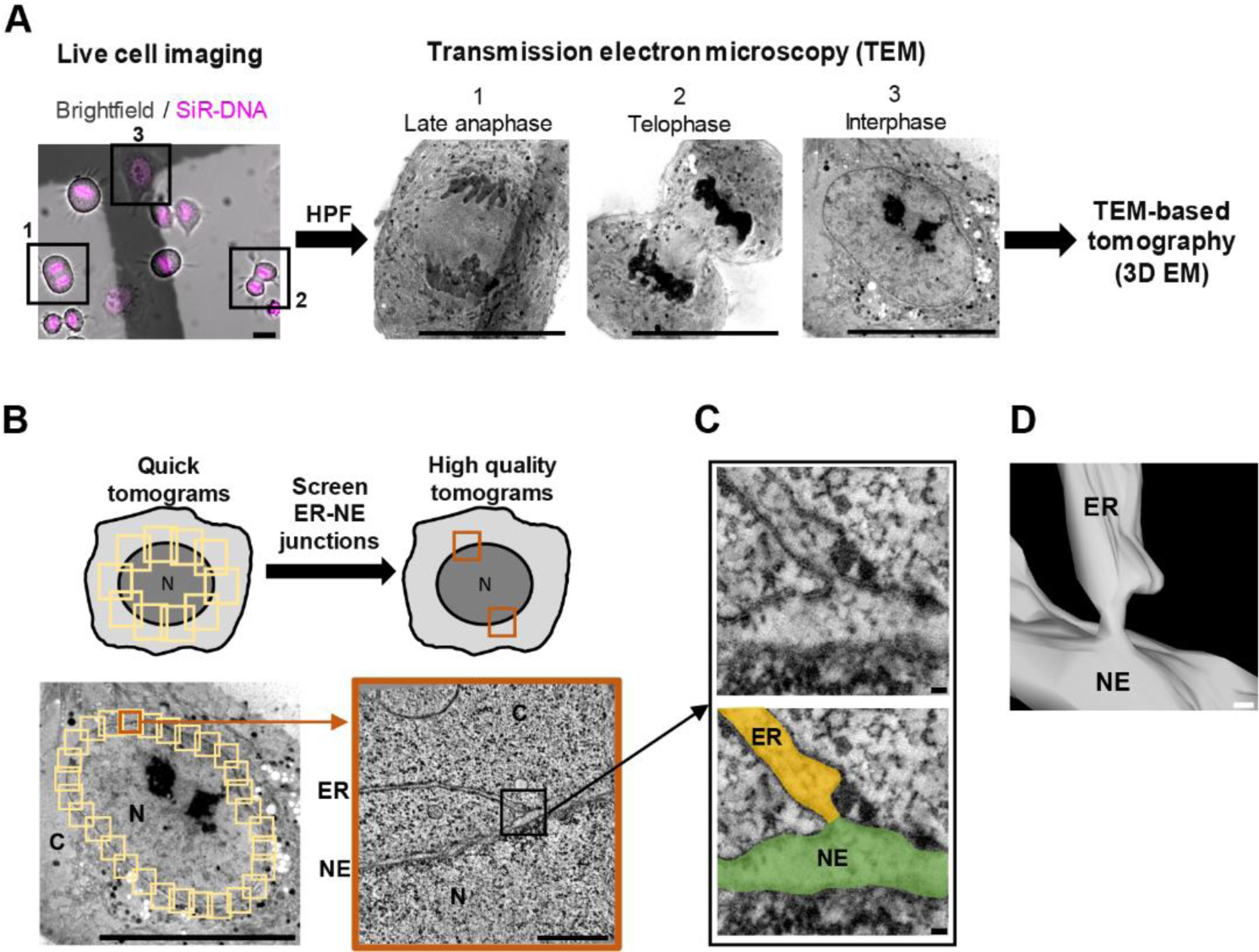
Method to systematically study the 3D ultrastructure of ER–NE junctions along the cell cycle. **A,** Correlative live cell imaging with electron microscopy to image the same cells using both imaging modalities. Cell-cycle progression of HeLa cells was monitored by live imaging in the bright-field and far-red (SiR-DNA) channels. Cells were high-pressure frozen (HPF) in late anaphase, telophase or interphase, and relocated under a transmission electron microscope (TEM) for tomography. Scale bars: 20 μm. **B,** Tomography-based screen to search for ER–NE junctions in an untargeted manner. Tomograms covering the entire NE were acquired quickly, and then high-quality tomograms were taken at regions with potential ER–NE junctions. Scale bars: 20 μm (overview), 500 nm (inset). **C,** An electron tomographic slice of an ER–NE junction. The membrane of the ER and the outer nuclear membrane are clearly visible in the high-quality tomogram. Scale bars: 20 nm. **D,** 3D mesh of the ER–NE junction shown in C. Scale bar: 20 nm. ER: endoplasmic reticulum, NE: nuclear envelope, C: cytoplasm, N: nucleus.

We developed a two-step tomography-based screen consisting of ‘quick’ and ‘high quality’ tomograms to identify ER–NE junctions in the cells in our samples (Figure 1B). The quick tomograms covered the entire NE and aimed to minimize the acquisition time of the tilt series while allowing sufficient image quality to identify regions with the ER and the NE in close proximity (Figure 1B). At these regions, high-quality tomograms were acquired to visualize the 3D ultrastructure of ER–NE junctions. The high-quality tomograms clearly resolved the lipid bilayer of the ER membrane continuous with the ONM of the NE (Figure 1C,D, Supplementary movie 1).

### ER–NE junctions are narrow and constricted in interphase

We characterized the ultrastructure of ER–NE junctions in interphase cells. High-quality tomograms revealed two distinct types of junction: those with a lumen (i.e. the ER/NE lumen was clearly visible at the interface) and those without a lumen (i.e. no lumen was visible at the interface, even though the membrane of the ER and the ONM appeared continuous) (Figure 2A, Supplementary Figure 1A,B). We also saw examples in which the ER membrane was juxtaposed to the ONM or connected to it by filament-like densities (Supplementary Figure 1C,D) that resemble the reported ‘contact sites’ between the ER and other intracellular organelles (Cai et al., 2022; Wozny et al., 2023). In this study, we focused on the ER–NE junctions in which the membranes of the ER and ONM appeared continuous.

**Figure 2.**
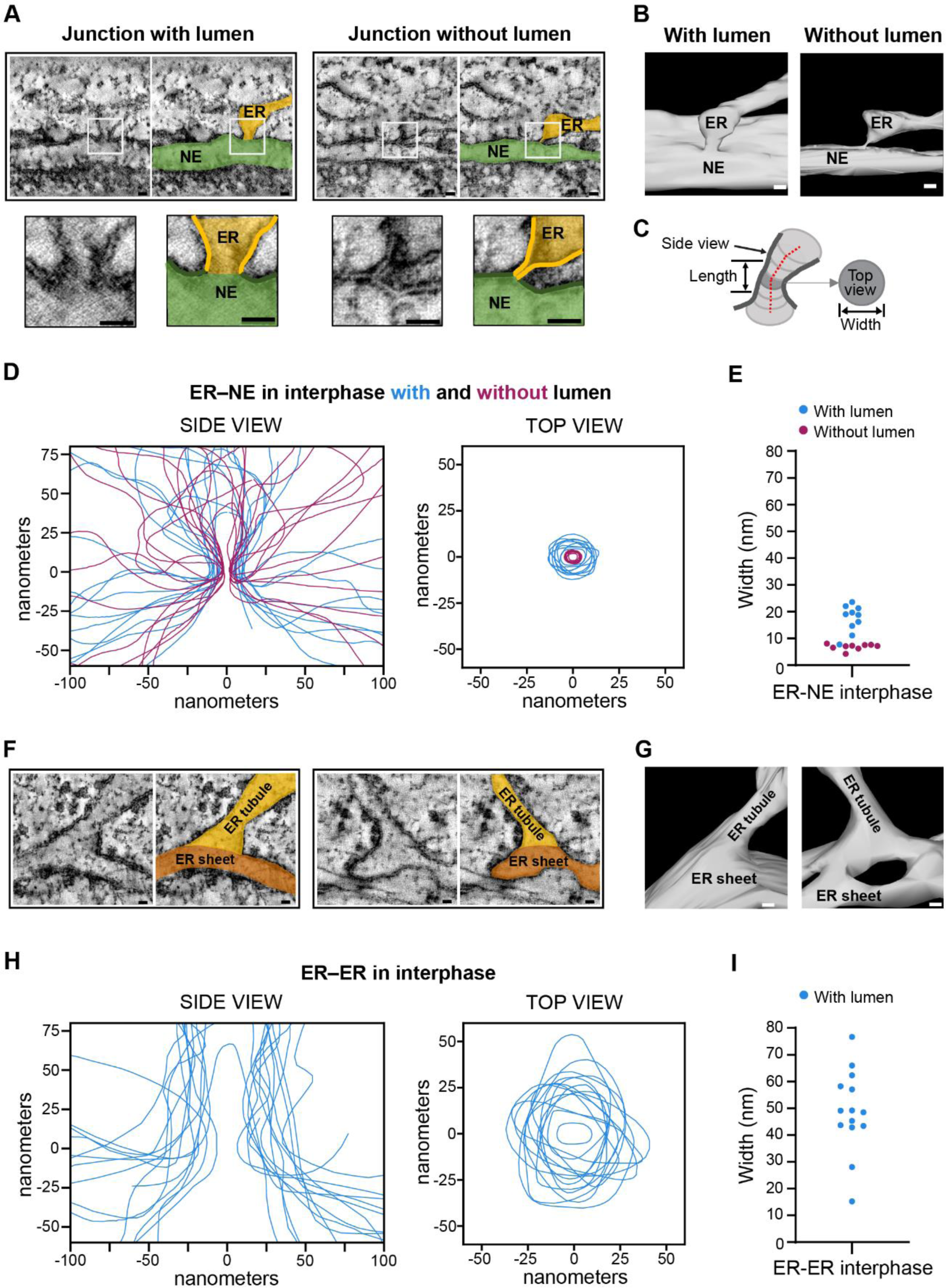
The membranous junctions that link the ER to the NE in interphase are narrow, constricted, and distinct from the junctions within the ER. **A**, Electron tomographic slices of ER–NE junctions with and without a lumen at their sagittal planes. For each junction, an enlargement of the region containing the junction is displayed below. The left image shows raw EM data; the right one, the EM data on which the ER and the NE are coloured in orange or green, respectively. Lipid bilayers are highlighted in bold lines in the insets. **B,** 3D meshes of the ER–NE junctions shown in **A**. **C,** The 3D mesh analysis allowed to obtain side and top view profiles, and the junction width and length. **D,** Membrane profiles of the side and top views of ER–NE junctions found in interphase cells. The profiles were drawn across the centre of the lipid bilayers. The side view profiles were overlaid vertically along the axis of the ER-NE junction. The top view profiles were aligned at their centroid. The profiles for the junctions with and without lumen are colour-coded in blue and magenta, respectively. n = 19 junctions from 9 cells. **E,** Width of the constricted neck of ER–NE junctions in interphase. n = 19, **F,** Electron tomographic slices of ER–ER junctions (in which ER tubules are fused perpendicularly to ER sheets) at their sagittal planes. The left image shows raw EM data; the right one, the EM data on which ER tubules and ER sheets are coloured in orange or brown, respectively. **G,** 3D meshes of the ER tubule-sheet junctions shown in **F**. **H,** Membrane profiles overlaying the side and top views of the ER tubule-sheet junctions found in interphase cells. The side and top view profiles were overlaid as in **D**. n = 14 junctions from 4 cells. **I,** Width of ER tubule-sheet junctions. n = 14. Scale bars: 20 nm.

To investigate in more detail the morphology of ER–NE junctions, we traced the ER/NE membranes in tomographic slices to generate a 3D model composed of meshes (Figure 2B). The 3D meshes allowed us to see the sagittal (side view) and transversal (top view) planes of the junctions, and to measure the width and length of the junction neck (Figure 2C, Supplementary Figure 2A–C). The sagittal membrane profiles revealed that the ER–NE junctions with and without lumen both had an hourglass shape with a constricted neck (n = 19 ER–NE junctions from 9 cells) (Figure 2D). Top view profiles showed that the neck was 17.4 ± 5.0 nm for junctions with a lumen and 6.9 ± 1.1 nm for junctions without a lumen (Figure 2E) (mean ± s.d., n = 10 junctions with a lumen, n = 9 junctions without a lumen). The width of junctions without a lumen corresponds to the thickness of a lipid bilayer, as the membrane profiles were drawn across the middle of the lipid bilayers. No ER–NE junctions larger than 24 nm in diameter were observed. The average length of the constricted neck was 10.4 ± 5.4 nm for junctions with a lumen and 4.9 ± 1.5 nm for junctions without a lumen (Supplementary Figure 2D–F). In addition, we noticed that the width of the perinuclear space below the junctions was larger than in other regions of the NE (Supplementary Figure 3). In short, our ultrastructural analysis demonstrated that ER–NE junctions at the interface between the ER and the NE have a pronounced hourglass shape with a constriction of 7–20 nm wide and a length of 4–10 nm.

To investigate whether the constricted morphology of ER–NE junctions is also seen in ER–ER junctions, we analysed the 3D morphology of ER–ER junctions in the same interphase cells in which the ER–NE junctions were observed above. We analysed specifically junctions that had a similar topology to ER–NE junctions, i.e. an ER tubule fused perpendicularly to an ER sheet. In contrast to ER–NE junctions, most ER–ER junctions had a wide cone-shaped base at the junction interface and much less marked membrane constrictions (Figure 2F–H) (n = 14 ER–ER junctions from 4 cells). All the ER–ER junctions had a lumen. The mean width of ER–ER junctions was significantly greater and more variable than that of ER–NE junctions (Figure 2I, Supplementary Figure 2G) (p-value <0.0001; Mann-Whitney test). Altogether, our ultrastructural analysis of interphase cells shows that ER–NE junctions have a highly curved and constricted hourglass shape that is distinct from the broad ER–ER junctions.

### ER–NE junctions become constricted in telophase

In metazoan cells that undergo open mitosis, the NE breaks down at the onset of mitosis and is absorbed in the ER (Ungricht & Kutay, 2017). The NE reforms again from membranes of the ER towards the end of mitosis in late anaphase and early telophase (Ungricht & Kutay, 2017). To see how and when ER–NE junctions reform during this process, we analysed the ultrastructure of ER–NE junctions when the NE is reforming. In late anaphase cells (4–6 minutes after the onset of anaphase), the ER and the NE are indistinguishable because the ER membranes just start contacting the surface of chromatin, presumably initiating NE reformation (Figure 3A–C, Supplementary Figure 4A–C). The junctions at these ER membranes touching chromatin were not constricted and had a similar morphology to ER–ER junctions in interphase. Consequently, the junctions at the ER that contact the chromatin in late anaphase have not yet adopted the specialised morphology typical of ER–NE junctions in interphase.

**Figure 3.**
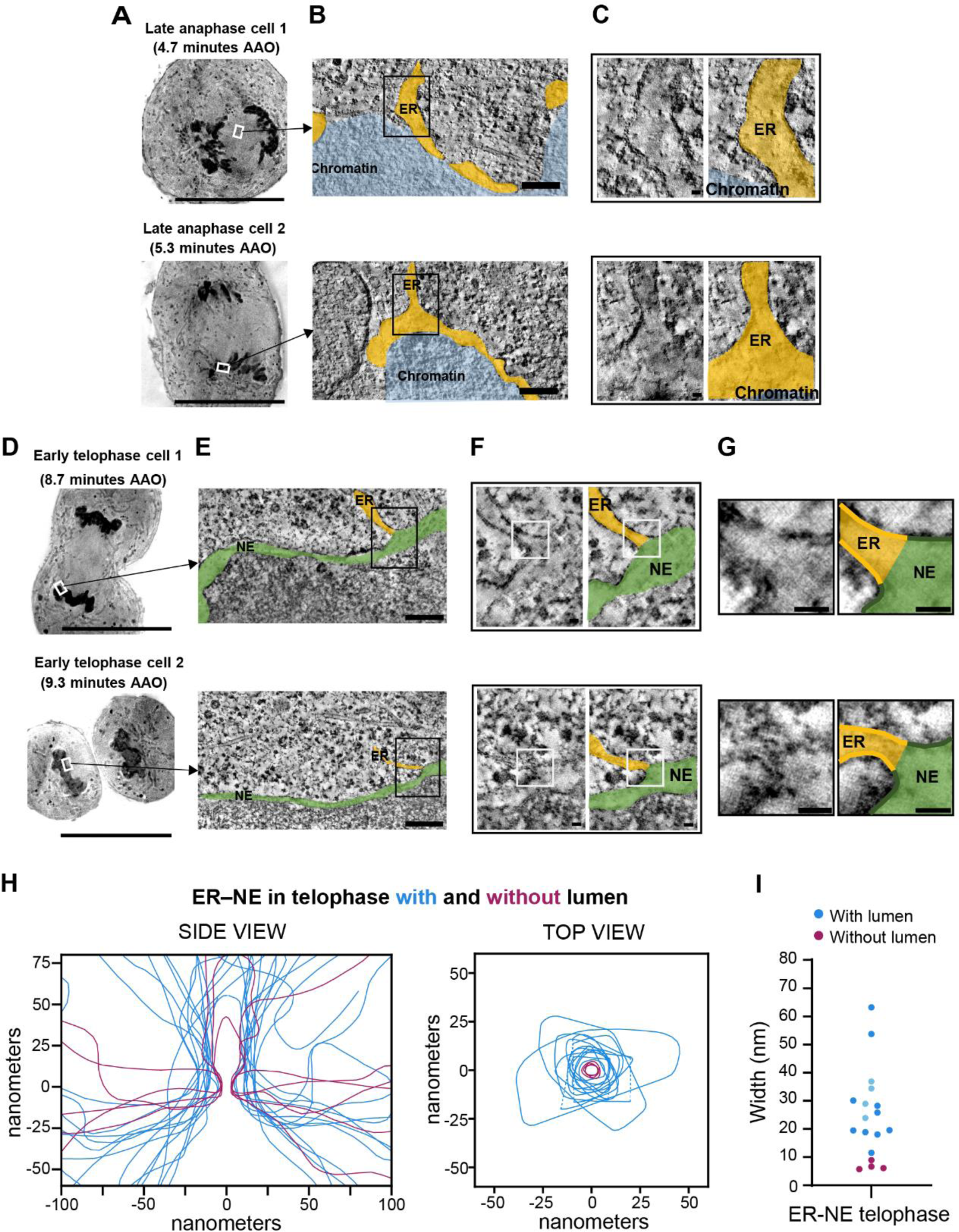
ER–NE junctions become progressively constricted during mitotic exit. **A**, 2D EM micrographs of two different cells in late anaphase. AAO: After Anaphase Onset. **B,** Tomographic slices of cells in late anaphase showing that the ER (orange) starts to contact the chromatin (blue), but the NE is still indistinguishable. **C,** Enlargement of the regions indicated in **B**. For each junction, the left image shows raw EM data; the right one, the EM data on which the ER and the chromatin are coloured in orange and blue, respectively. **D,** 2D EM micrographs of two different cells in early telophase. **E,** Tomographic slices showing that ER–NE junctions are present on the NE reformed on the chromatin in early telophase. **F,** Enlargement of the regions indicated in **E**. **G,** Enlargement of the regions indicated in **F**. For each junction in **F** and **G**, the left image shows raw EM data; the right one, the EM data on which the ER and the NE are coloured in orange and green, respectively. Lipid bilayers are highlighted in bold lines in **G**. **H,** Membrane profiles overlaying the side and top views of the ER–NE junctions found in early telophase cells. The side and top view profiles were drawn as in Fig. 2D. n = 18 junctions from 2 cells. In 4 junctions, the entire top views could not be traced in the EM tomograms as they were truncated at the edge of the tomograms. For these 4 junctions, dashed lines indicate the truncated edge of the top view profiles. **I,** Width of ER–NE junctions in early telophase. n = 18. Dots in light blue indicate the 4 truncated top view profiles, whose width is underestimated. Scale bars for **A, D**: 20 μm; Scale bars for **B**, **E**: 200 nm; Scale bars for **C, F, G**: 20 µm.

To investigate when the constricted structure of ER–NE junctions begins to appear, we looked in early telophase cells (8–10 minutes after the onset of anaphase). In telophase, the NE was clearly distinct from the ER because the membrane was in direct physical contact with the chromatin and covered almost all the surface of the daughter nuclei (Figures 3D–G, Supplementary Figure 4D–G). Ultrastructural analysis revealed that most of ER–NE junctions in telophase start to become constricted (Figure 3H) and distinct from ER–ER junctions (Figure 2H). The average width of top view profiles was 26.2 ± 7.9 nm (mean ± s.d., n = 18 ER–NE junctions from 2 cells) (Figure 3I), which is between that of ER–NE and ER–ER junctions in interphase (Supplementary Figure 2G), indicating that the membrane constriction is not complete. These data suggest that the constriction of ER–NE junctions starts in early telophase and undergoes further maturation to form fully specialized junctions.

### The number of ER–NE junctions per cell increases from telophase to interphase

To examine whether the abundance of ER–NE junctions changes from telophase to interphase, we quantified the NE surface area in the EM tomograms previously screened for junctions (Figure 4A,B) in order to calculate the frequency of ER–NE junctions in telophase and interphase. In most junctions we could distinguish clearly the ER membrane continuous with the ONM (as in Figure 2A, Supplementary Figure 1A,B). A few junctions were categorized as ambiguous when the ER/NE membranes could not be traced with enough confidence. These ambiguous junctions had only a small effect on our calculations because they were relatively few and similar in frequency in telophase and interphase cells (Figure 4C). The frequency of all ER–NE junctions was ∼0.18 μm^-2^ in telophase (21 junctions in a screened NE surface area of 112 μm^2^ from 2 cells) and ∼0.13 μm^-2^ in interphase (23 junctions in a screened NE surface area of 178 μm^2^ from 9 cells) (Figure 4C). These densities were much lower than those of ER–ER junctions or other NE-specific structures like nuclear pore complexes, which are 50–100 times more abundant (Tikhomirova et al., 2022; Otsuka et al., 2018).

**Figure 4.**
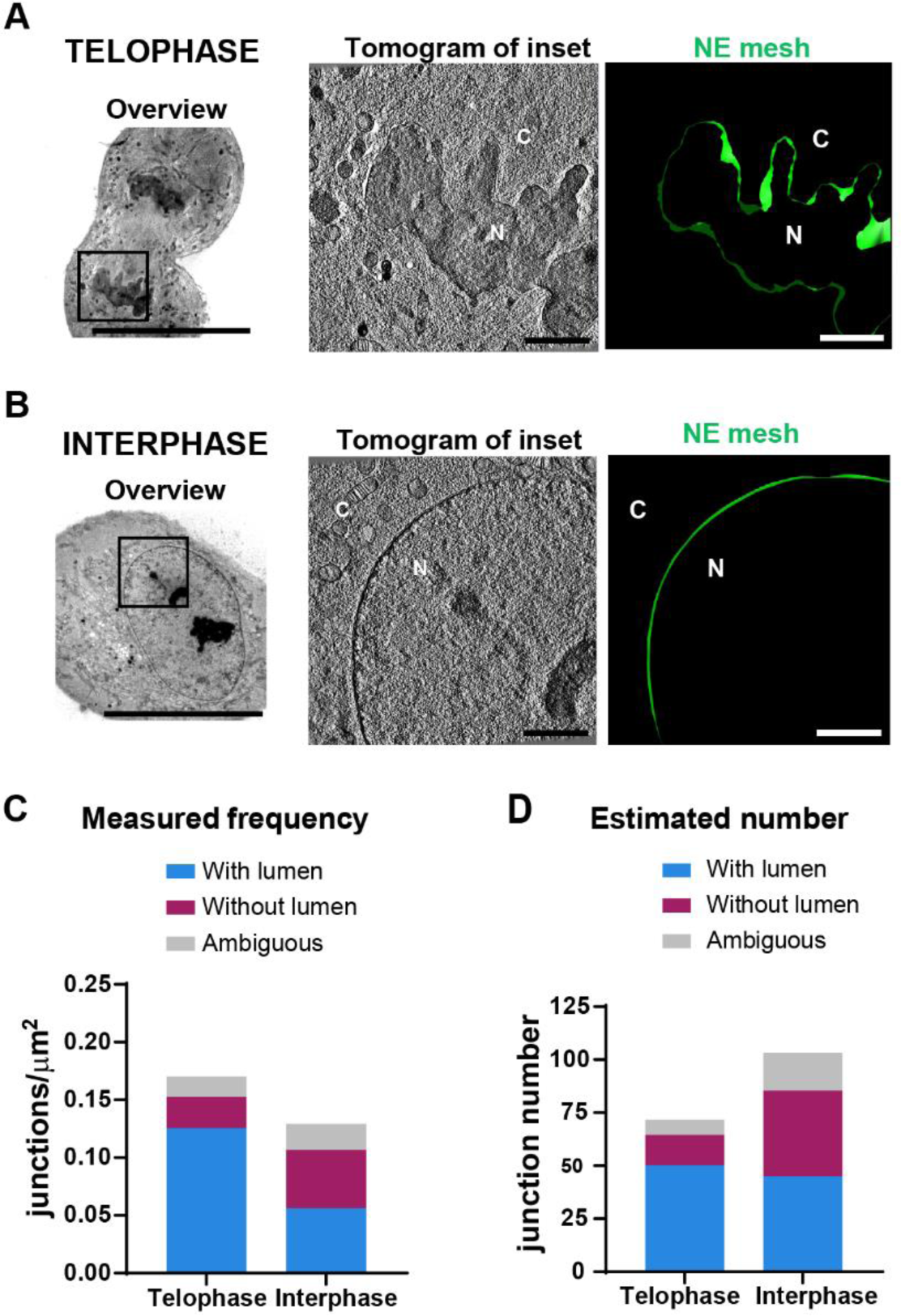
The amount of ER–NE junctions changes slightly from early telophase to interphase. **A,B** Making 3D meshes of the NE for quantifying its surface area observed by EM tomography in early telophase (**A**) and interphase (**B**) cells. Left: 2D-EM micrograph of each overview image; Middle: a tomographic slice of the region indicated in each overview image; Right: a 3D mesh of the NE obtained from the tomogram shown in the middle panel. C: cytoplasm, N: nucleus. Scale bars for overview: 20 μm; for inset: 200 nm. **C,** Frequencies of the different types of ER–NE junctions in early telophase and in interphase cells calculated by dividing the number of junctions by the surface area quantified in **A** and in **B**. **D,** Estimated total number of ER–NE junctions in average nuclei of early telophase and interphase cells, calculated by multiplying the frequencies in **C** with the total NE surface of early telophase and interphase nuclei measured in a previous study (400 μm^2^ for early telophase and 800 μm^2^ for interphase) (Otsuka et al., 2016).

We next asked if the total number of ER–NE junctions per nuclei increases in response to the nuclear growth from telophase to interphase. A previous study reported that HeLa cells have a total NE surface area of about 350–450 μm^2^ in telophase (at 8–10 minutes after the onset of anaphase), and 700–900 μm^2^ in interphase (Otsuka et al., 2016). Using these reported values of NE surface area and assuming a constant frequency of junctions around the whole nucleus, we estimated that a typical nucleus has approximately 70 junctions in telophase and 100 junctions in interphase (Figure 4D). The number of junctions with a continuous lumen was similar in telophase and interphase cells, whereas the number junctions without a lumen was greater in interphase (Figure 4D). These data show that the number of ER–NE junctions increases slightly from telophase to interphase, although their abundance remains considerably lower than that of ER–ER junctions or nuclear pores at both cell cycle stages.

### The constricted morphology of ER–NE junctions is observed in different mammalian cells, but not in budding yeast

To determine if the constricted morphology of ER–NE junctions in interphase HeLa cells is common to other mammalian cells, we used whole-cell EM datasets publicly available in OpenOrganelle (Heinrich et al., 2021; Xu et al., 2021) to quantify ER–NE and ER–ER junctions in mouse pancreatic islet, HeLa, and human macrophage cells. In these datasets, entire cells were imaged by focused ion beam (FIB) scanning EM (FIB-SEM) at a voxel size of around 4 nm, which allowed us to investigate ER–NE junctions around the whole NE (Supplementary Figure 5). Although the spatial resolution of the FIB-SEM datasets is ten times lower than our EM tomograms, it was sufficient to identify and quantify the overall width of potential ER–NE junctions (Figure 5A–E). To study ER–NE junctions in an unbiased manner, we took a stereology-based approach. The regions of interest were sampled in 1 μm radius spheres distributed uniformly over the entire nuclear surface, and those regions were inspected for potential ER–NE junctions (Supplementary Figure 5). This unbiased inspection of the FIB-SEM datasets revealed that the width of ER–NE junctions in the pancreatic islet cells (Figure 5A), HeLa (Figure 5B), and macrophage (Figure 5C) were significantly smaller than most ER–ER junctions (Figure 5F), which is consistent with the ultrastructural observation in our high-quality EM tomograms (Figure 2). Due to the limited spatial resolution of the FIB-SEM data, it is difficult to distinguish the ER–NE junctions with continuous membranes from the ER–NE ‘contact sites’ that we identified by tomography (Figure 2A and Supplementary Figure 1). Therefore, the ER–NE junctions that we analysed in the FIB-SEM datasets may contain ER membranes that are in close proximity to, but not necessarily fused with, the ONM. Nevertheless, we rarely found ER–NE junctions that were not constricted. This analysis of FIB-SEM images confirms the hourglass morphology that distinguishes ER–NE from ER–ER junctions as seen in our EM tomograms of HeLa cells, and extends our findings to two other mammalian cell types.

**Figure 5.**
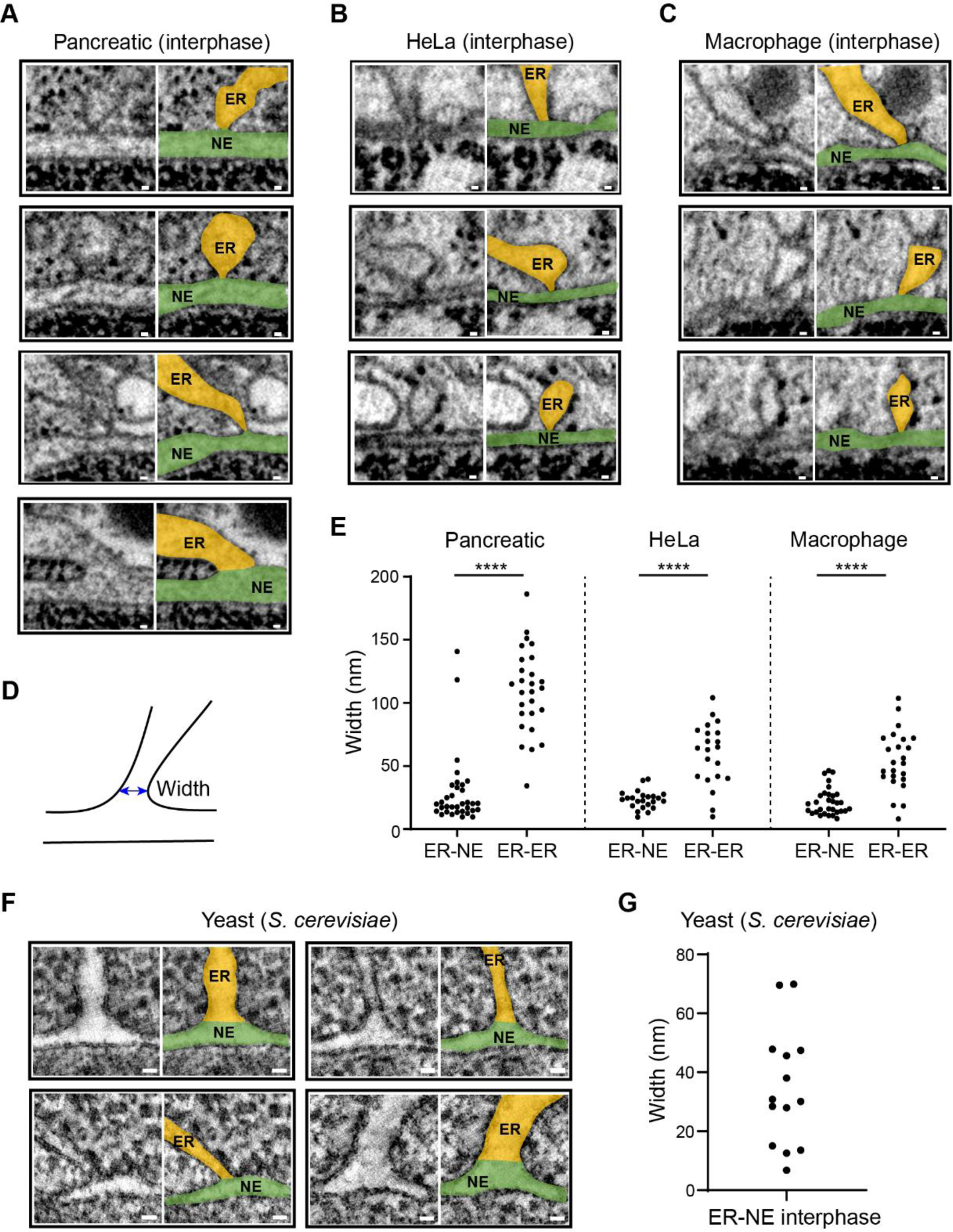
ER–NE junctions are constricted in several mammalian cell lines, but not in budding yeast. **A–C**, ER–NE junctions in FIB-SEM images of a mouse pancreatic islet cell (pancreatic) (**A**), a HeLa cell (**B**), and a human macrophage (**C**). For each junction, the left image shows raw EM data; the right one, the EM data on which the ER and the NE are coloured in orange or green, respectively. The images in the fourth row in **A** show one of the two junctions in the pancreatic cell with a width above 100 nm. **D,** Scheme depicting how the width of junctions was measured using a 2-point line (blue). **E,** Widths of ER–NE and ER–ER junctions in a mouse pancreatic islet cell, a HeLa cell, and a macrophage. n = 34 ER–NE and 27 ER–ER junctions for a mouse pancreatic islet cell, n = 23 ER–NE and 21 ER–ER junctions for a HeLa cell, and n = 31 ER–NE and 24 ER–ER junctions for a macrophage. ****p-value < 0.0001; two-tailed Mann–Whitney test. **F,** Tomographic slices of ER–NE junctions of budding yeast (*Saccharomyces cerevisiae*) cells at their sagittal planes. **G,** Widths of ER–NE junctions measured in the yeast cells. n = 14 junctions from 8 cells. Scale bars for **A–C, F**: 20 nm.

To investigate whether hourglass-shaped ER–NE junctions are conserved among eukaryotes, we visualised ER–NE junctions in high-pressure frozen *S. cerevisiae* cells by tomography (Figure 5F). In agreement with a previous study (West et al., 2011), we observed junctions between the NE and ER cisternae and tubules (Figure 5F). Most ER– NE junctions had a wide cone-shaped base at the junction interface and no particular constriction (Figure 5F). Quantification of the junction widths showed that ER–NE junctions in budding yeast are variable (Figure 5G). We conclude that ER–NE junctions are generally not constricted in budding yeast; the highly constricted hourglass morphology of ER–NE junctions is a specialized feature of mammalian cells.

## DISCUSSION

Whereas the molecular mechanisms that govern ER–organelle contact sites and their functions have been studied extensively (Rossini et al., 2021), the structure and the physiological role of the junctions connecting the ER to the NE remain poorly understood. Our study using correlative light and 3D electron microscopy reveals the native ultrastructure of ER–NE junctions in high-pressure frozen mammalian cells at unprecedented spatial resolution. In interphase cells, the ER forms hourglass-shaped junctions with the ONM of the NE that are highly curved with a narrow constriction (7– 20 nm in diameter). These ER–NE junctions are clearly distinct from the broad ER–ER junctions. The width of the perinuclear space below the junctions was larger than in other regions of the NE (Supplementary Figure 3), suggesting that LINC (linker of nucleoskeleton and cytoskeleton) complexes that typically maintain a NE width of 30–50 nm do not form below ER–NE junctions (Cain & Starr, 2015). During reassembly of the NE in late anaphase the junctions resemble ER–ER junctions, and they start to become constricted in early telophase cells (Figure 6A). This indicates that the formation of the hourglass shape of ER–NE junctions is initiated rapidly after mitosis and presumably maintained for the rest of the cell cycle. Our study is in agreement with the observations made by Craig and Staehelin over three decades ago that plant cells have constricted ER– NE junctions when they are high-pressure frozen (Craig and Staehelin, 1988). We show at high resolution and in 3D that the constricted morphology of ER–NE junctions is a common feature in mammalian cells. We also confirm a previous finding that the ER– NE junctions of budding yeast have a non-constricted shape (West et al., 2011). A cryo-EM tomogram of a single ER–NE junction in *Chlamydomonas* also showed no constriction of the junction (Albert et al., 2017). Thus, our work suggests that the hourglass morphology of ER–NE junctions emerged only in higher eukaryotes, and evidence from several other species would be necessary to consolidate it.

**Figure 6.**
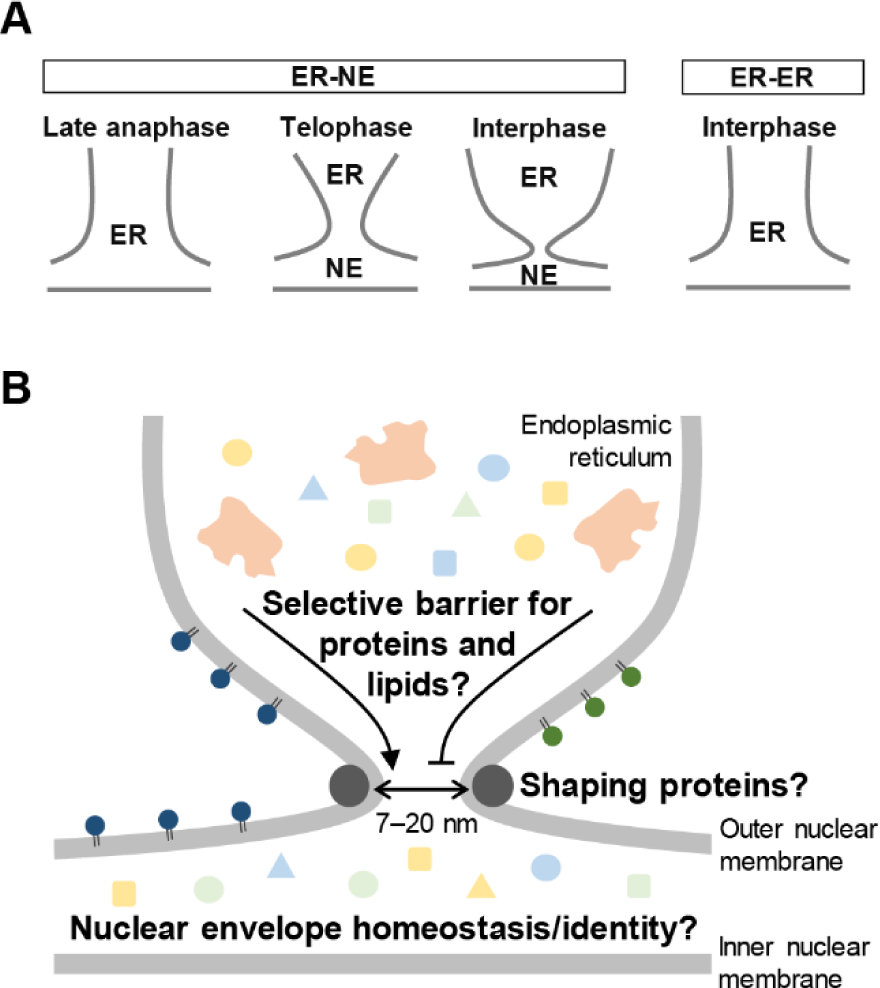
Model for the formation and function of ER–NE junctions in higher eukaryotes. **A**, ER–NE junctions mature progressively after telophase into a constricted hourglass-shaped morphology that is distinct from the broad junctions within the ER. **B,** Implications and potential functions associated with the constricted morphology of ER– NE junctions in interphase.

What biophysical and molecular mechanisms might generate the high curvature of ER–NE junctions? Evidence from *in vitro* systems composed entirely of lipids indicate that the generation and stabilization of such a morphology is energetically unfavourable in the absence of external forces (Lipowsky, 2022). We hypothesize the existence of membrane remodelling proteins that are localized at the neck of ER–NE junctions to maintain their constricted shape (Figure 6B). Proteins known to form and stabilize junctions in the ER, including Atlastins and Lunapark (Wang et al., 2016; Pawar et al., 2017), might be involved in generating and stabilizing ER–NE junctions. However, given that ER–NE junctions are clearly distinguished from ER–ER junctions in morphology and abundance, additional proteins are presumably required to differentially remodel ER– NE junctions. Our discovery that ER–NE junctions become constricted in telophase suggests that special remodelling proteins might be recruited to the NE as early as telophase. These remodelling proteins might be expressed predominantly or exclusively in higher eukaryotes, as ER–NE junctions are not highly constricted in yeast and *Chlamydomonas*. Unfortunately, our EM tomograms precluded the detection of consistent protein densities in the lumen or around the neck of ER–NE junctions that might correspond to large structural protein complexes. Future studies combining high-resolution imaging with molecular perturbations of the system will be necessary to identify potential regulators responsible for shaping and stabilizing ER–NE junctions.

How might ER–NE junctions form *de novo* during interphase? We found that the number of ER–NE junctions whose lumens are clearly connected does not change from telophase to interphase, whereas the number of junctions whose lumens do not connect increases 4-fold (Figure 4D). Thus, we postulate two hypotheses for the biogenesis of ER–NE junctions. One hypothesis is that the junctions with a lumen are maintained throughout the cell cycle, whereas the junctions without a lumen arise *de novo* during interphase. The other hypothesis is that both types of ER–NE junctions are constantly remodelled (formed and degraded) during the cell cycle, like ER–ER junctions that form and break constantly during interphase (Nixon-Abell et al., 2016; Takakura et al., 2017). In either case, at least a small fraction of ER–NE junctions would form *de novo* from telophase to interphase. Regarding *de novo* formation of ER–NE junctions, we envision two scenarios. In one scenario, the NE would extend membrane protrusions into the cytoplasm. However, we observed no examples of protruding ONM in our EM images. Thus, we favour a second scenario in which ER–NE junctions are generated from ER tubules that contact and eventually fuse with the ONM. In our tomograms we found cases in which the ER membrane was juxtaposed (but not fused) to the ONM (Supplementary Figure 1C,D). These contact sites may be assembly intermediates, and certain proteins may be recruited to mediate membrane fusion.

Given the constricted ultrastructure of ER–NE junctions, it is tempting to speculate that ER–NE junctions may act as a selective barrier for the transport of lipids, large protein complexes, and protein aggregates between the ER and the NE (Figure 6B). A previous study showed that phosphatidylserine (one of the major acidic phospholipids in eukaryotic cells) is over 10-fold enriched in the ER membrane when compared with the ONM despite the continuous nature of the ER and NE, and this phosphatidylserine enrichment occurs in mammalian cells and not in budding yeast (Tsuji et al., 2019). Since phosphatidylserine is a cylindrical-shaped lipid that favours the formation of flat membranes (Peeters et al., 2022), ER–NE junctions with highly-curved membranes may exclude phosphatidylserine and prevent its diffusion into the ONM. As NE lipid composition affects NE protein stability (Lee et al., 2023), ER–NE junctions may thus play a key role in NE lipid/protein homeostasis, as proposed previously (Bahmanyar & Schlieker, 2020). The average width of ER–NE junctions of 7–20 nm is larger than the diameter of most proteins (∼5 nm), but small enough to restrict the diffusion of large multimeric complexes and protein aggregates between the lumens of the ER and NE. Interestingly, Apomucin, which forms oligomers of 500–1000 kDa (Eckhardt et al., 1987), localises predominantly in the ER lumen rather than the NE lumen in mammalian cells (Deschuyteneer et al., 1988). ER–NE junctions may act as a size-selective gate that prevents large protein complexes and misfolded proteins to enter the perinuclear space (Sun & Brodsky, 2019). In this way, the constricted junctions may protect the NE from the potential toxicity of misfolded/aggregated proteins. The two types of ER–NE junctions we observed – with and without a luminal connection – raise the possibility that ER–NE junctions can change configuration. The membrane contact sites between the ER and other organelles such as mitochondria, endosome, and plasma membrane respond to various environmental cues including nutrient stress, ER stress and calcium stimulation (Donahue et al., 2022; Jang et al., 2022). Similarly, the membrane junctions between the ER and the NE might change their morphology in response to environmental stimuli. It would be interesting to examine the dynamic nature of the junctions under physiological stresses.

Newly synthesised NE proteins that cross ER–NE junctions play crucial roles in gene expression, nuclear organization, and nuclear pore biogenesis, as well as in development and disease (Gauthier and Comaills, 2021). ER–NE junctions may contribute to these key nuclear functions by acting as selective barriers for proteins and lipids. Dysregulated nuclear size, mislocalised NE proteins, and dysfunction of ER-shaping proteins are associated with many diseases including cancer and neurodegenerative disorders (Jevtic and Levy, 2014; Öztürk et al., 2020; Rose et al., 2022). Few physiological processes have been linked to ER–NE junctions, perhaps due to the lack of studies focusing on this particular type of junction. Our finding that ER– NE junctions have a distinct hourglass morphology in mammalian cells provides a basis for future molecular and functional studies in the context of physiology and disease.

## METHODS

### Cell culture

Wild-type HeLa Kyoto cells (RRID: CVCL_1922) were incubated in Dulbecco’s Modified Eagle Media (DMEM with low glucose, Sigma, D6046-500ML) supplemented with 10% fetal calf serum (FCS, Sigma, F7524-500ML) and 1% Penicillin-Streptomycin (Pen/Strep, Sigma, P4333-100ML). Cells tested negative for mycoplasma infection using a mycoplasma detection kit (Biological Industries, 20-700-20).

### Cell seeding and cell cycle synchronization for correlative light and electron microscopy

Carbon-coated sapphire discs (D6mm, Leica Microsystems) were assembled on 1-well cups for SampLink chambers (Leica Microsystems). The assembled SamplLink chambers were washed in 70% ethanol, exposed to UV light for 20–30 minutes, and dried overnight. Before seeding the HeLa cells, the sapphire discs were coated with 0.1 mg/ml poly-L-lysine hydrobromide (Sigma, P1274) for 1–1.5 hours at 37 °C. Few hours after seeding the cells, when most cells had adhered to the sapphire disc, the media was replaced with 2 mM thymidine (Sigma, T9250-1G) in DMEM+FCS+Pen/Strep media overnight. On the next day, samples were washed twice in Dulbecco’s Phosphate Buffered saline (DPBS, Sigma, D8537) and incubated in fresh DMEM+FCS+Pen/Strep media to release cells from the first thymidine block. Six hours later, samples were washed in PBS and incubated in 2 mM thymidine in DMEM+FCS+Pen/Strep overnight. On the following day, samples were washed twice in PBS and incubated in fresh DMEM+FCS+Pen/Strep media to release cells from the second thymidine block.

### Live cell imaging and high-pressure freezing

Eight hours after the release from the second thymidine block, the media was replaced with imaging media (DMEM without Riboflavin and Phenol Red, Thermo Fisher Scientific, Gibco 041-96205M, containing 10% FBS, 1% Pen/Strep, and 50 nM SiR-DNA, Spirochrome, SC007). An hour later, samples were transported to the light microscope using the Baker Ruskinn OxyGenie transportable incubator (Leica Microsystems) at 37 °C equipped with gas cylinders (Catalina Cylinders) filled with 5% CO_2_ in a synthetic atmosphere of 20% O_2_ and 80% N_2_. Cells in the SampLink chambers were monitored in the light microscope THUNDER Imager Nano (Leica Microsystems) coupled with an incubator at 37 °C. Time-lapses of regions enriched in metaphase cells were acquired every 20 seconds in the bright-field and far-red (SiR-DNA) channels using a 20x 0.40 NA DRY NPLAN Epi objective (Leica Microsystems). When most dividing cells in the field of view reached late anaphase and telophase, samples were high-pressure frozen using the High Pressure Freezer Leica EM ICE equipped with the Coral Life workflow (Leica Microsystems). It took 1.0‒1.5 minutes from the last time-lapse imaging until the high-pressure freezing, and the time lag was recorded to precisely determine the duration after anaphase onset. Cells were frozen in DMEM+FCS+Pen/Strep supplemented with 20% FICOL-PM400 (Sigma, F4375-25G) as a cryoprotectant, which was added to the samples few seconds before freezing. The lid carriers (Leica Microsystems) had a cavity depth of ∼25 μm, and they had been in contact with 1-hexadecene (Merck, 8.22061.0100). After high-pressure freezing, sapphire discs were stored in liquid nitrogen until the subsequent sample preparation steps.

### Freeze substitution and resin embedding

Freeze substitution was performed in a Leica EM AFS-2 freeze substitution unit (Leica Microsystems) as described previously (Bragulat-Teixidor et al., 2022). Briefly, the samples were substituted in 0.1% uranyl acetate (UA), 2% Osmium tetroxide (Electron Microscopy Sciences, 19134) and 5% H_2_O in acetone (PanReac AppliChem, ITW reagents, 141007.1211) following this temperature ramp: -90 °C to -80 °C for 10 hours, -80 °C to -30 °C for 10 hours, -30 °C for 4 hours, -30 °C to 0 °C for 6 hours, 0 °C to 20 °C for 4 hours, 20 °C for 5–6 hours. Afterwards, samples were washed three times in pure acetone for at least 10 minutes each, and subsequently infiltrated with Agar 100 Epoxy resin (Agar Scientific). The resin infiltration was done progressively at room temperature with increasing concentrations of resin in acetone (3:1 for 2–3 hours, 1:1 for 2–3 hours, and 1:3 overnight). Infiltration with pure resin was done at room temperature for at least 5 hours. Resin was polymerized at 60 °C for 72 hours.

### Sectioning, gold bead attachment and post-staining

Resin blocks were sectioned every 250 nm using a Diamond knife (Diatome) and an ultramicrotome Leica ultracut UCT (Leica Microsystems). Sections were collected on Cu/Pd slot grids (Agar Scientific, G2564PD) coated with a film of 1% formvar (Agar Scientific) in chloroform (Sigma Aldrich, 32211-1L-M). To allow better alignment of dual-axis tomography, 15-nm gold beads conjugated with Protein A (Cytodiagnostics, AC-15-05-10) were attached to both sides of the grids. To enhance membrane contrast, sections were post-stained at room temperature with 2% uranyl acetate in 70% methanol for 10 minutes and with 3% Reynold’s lead citrate in H_2_O (Delta Microscopies) for 7 minutes.

### Electron tomography

As ER–NE junctions are too tiny to be identified reliably on 2D micrographs, we developed a two-step tomography-based screen consisting of ‘quick’ and ‘high quality’ tomograms. The first step aimed at minimizing the acquisition time of the tilt series while allowing sufficient image quality to resolve the ER from NE membranes. These tilt series were acquired over a ± 20° tilt range with an angular increment of 2° at a pixel size of 0.567 nm. These tomograms were taken with slight overlaps to cover the entire NE in one or more sections per cell (Figure 1B). These tilt series allowed the reconstruction of ‘quick’ tomograms, which were screened manually for potential ER–NE junctions. In the second step of the screen, ‘high quality’ tomograms (typically dual axis, from tilt series with a ± 50° tilt range, an angular increment of 1°, and a pixel size of 0.451 nm) were taken at the regions where the ‘quick’ tomograms revealed potential ER–NE junctions.

We used a similar two-step tomography-based screen to search for ER–ER junctions with a similar topology to ER–NE junctions (i.e. an ER tubule contacting perpendicularly to an ER sheet) in the same cells where ER–NE junctions were analysed. We acquired ‘low magnification’ tomograms (from tilt series with a ± 50° tilt range, an angular increment of 1°, and a pixel size of 1.137 and 2.181 nm) and ‘high quality’ tomograms (typically dual axis, from tilt series with a ± 50° tilt range, an angular increment of 1°, and a pixel size of 0.451 nm). The ‘low magnification’ tomograms allowed to find ER–ER junctions in the cytoplasm; the ‘high quality’ tomograms, to image the junctions at a high magnification for a better visualization and quantification.

The tomograms were reconstructed from tilt series with the R-weighted backprojection method implemented in the IMOD software package (version 4.11.7) (Kremer et al., 1996). The tilt series were acquired using the Serial EM software (v3.x) under the transmission electron microscope (TEM) Tecnai G2 20 operated at 200 kV and equipped with an Eagle 4k HS CCD camera. The Stitching plugin (Preibisch et al., 2009) in FIJI (Schindelin et al., 2012) was used to efficiently relocate the tomograms in stitched overviews of the cells.

### Quantification of the frequencies of ER–NE junctions

The frequency of ER–NE junctions was calculated by dividing the total junction number by the total NE surface area screened for junctions in quick tomograms. The junctions were identified in high-quality tomograms as described previously. The NE area was measured from quick EM tomograms, taking into account the overlay among consecutive overlapping tomograms. For telophase, we calculated the NE from meshes obtained by manually tracing the NE using IMOD. In general, we traced the NE on three tomographic planes (at beginning, middle and end) and interpolated the NE traces along the tomographic axis. Since the NE is highly curving in telophase cells (Figure 4A), the NE interpolation sometimes mismatched the raw EM data. Whenever there was a mismatch, additional contours were drawn on the NE so that the mesh could represent the true NE. For interphase, the NE is flat and almost perpendicular to the viewing plane of the section (Figure 4B), so the NE area was estimated by multiplying the length of the NE in 2D overview images with the depth of the NE from the start to the end of the section. Concretely, we first identified regions with different tilts of the NE on 2D overviews of cells, and measured their length. Then, we measured the thickness of the NE in a quick tomogram of that region by a 2-point line in IMOD. Finally, the NE area was calculated by multiplying the length of a certain NE region with the depth of the NE in that same region. We confirmed the robustness of the method using three cells by comparing the NE surface area obtained in IMOD meshes (as done for telophase cells) versus the area obtained after multiplying the length with the depth of the NE on sections. The NE area values obtained by either method differed only up to ∼3 %, which confirms the reliability of this measurement.

### Measurement of the NE width below ER–NE junctions

The NE width was measured as a 2-point line spanning the perinuclear space below ER– NE junctions and at regions 200-500 nm away from ER–NE junctions and nuclear pore complexes (Supplementary Figure 3). The measurements were done in IMOD after orienting the tomograms in a way such that the NE would be horizontal and perpendicular to the viewing plane. Then, below junctions, the width was measured from the base of the junction (where the ONM would be present if the junction would be absent) to the closest point at the inner nuclear membrane (INM). Away from junctions, the width was measured from a point at the ONM to the nearest point at the INM. The distance away from junctions was measured twice for each junction (i.e. in two opposite directions away from the junction).

### Electron microscopy of yeast cells

Wild-type strains of budding yeast (*Saccharomyces cerevisiae*, BY4741, Euroscarf) cells were cultured at 30 °C in synthetic dextrose complete media. Asynchronous yeast cells were high-pressure frozen at the log phase and freeze substituted as described previously (Romanauska and Köhler, 2021). Sections with a thickness of 400 nm were cut using a Leica UCT ultramicrotome (Leica Microsystems). The grids were coated with 15-nm gold beads conjugated with Protein A on both sides, and the sections were post-stained with 2% UA in methanol and 3% Reynold’s lead citrate in H_2_O as described above. Cells were observed using the Serial EM software (v3.x) under the transmission electron microscope (TEM) Tecnai G2 20 operated at 200 kV and equipped with an Eagle 4k HS CCD camera. At the regions where ER was visible near the NE, dual axis tomograms were acquired typically over a -60° to +60° tilt range with an angular increment of 1° at a pixel size of 0.451 nm.

### Mesh generation

Meshes of ER–NE junctions were generated using the IMOD software after manually tracing the ER membrane, ONM and INM across the middle of their lipid bilayer every 2.25–4.51 nm tomographic slices. For meshes of ER–ER junctions, the membranes were traced every 9.02 nm tomographic slices.

### Side profiles of ER–NE and ER–ER junctions in tomograms

The side profiles were drawn by tracing manually the ER membrane and the ONM across the middle of the lipid bilayer after rotating the EM tomogram to the orientation of the sagittal plane of the junction in the Slicer mode of IMOD (Kremer et al., 1996; version 4.11.7).

### Cross-section analysis for determining top profiles, width and length of ER–NE and ER–ER junctions in tomograms

For each mesh, cross-sections were generated from the junction base at fixed intervals (typically 0.4 to 2.0 nm which varied for each example) along a centreline that spanned the neck of the junction using the NeuroMorph Analysis and Visualization Toolkit (Jorstad et al., 2018) in Blender (version 2.83). The cross-sections allowed to obtain the top profiles, and to analyse the width and length of junctions. The top profiles were drawn manually across the middle of the lipid bilayer at the neck of each junction after orienting the EM tomograms in the Slicer mode of IMOD (Kremer et al., 1996; version 4.11.7).

The orientation and position corresponded to the one of the cross-section with the minimum surface area within 25 nm away from the base of the junction. To identify the cross-section with the minimal surface area, the area of all cross-sections was measured using a custom script based on Python 3.9.5 using the libraries Trimesh 3.10.8 (Dawson-Haggerty et al., 2019) and Shapely 1.8.5 (Gillies et al., 2022). The width of junctions was defined as the average of the longest and shortest lengths of the bounding rectangle around the top profiles. The measurements were done using a custom-made script. The length of a junction L resulted from the sum of two shorter length measurements named L_1_ and L_2_, where L_1_ was the distance from the minimum area cross-section to the closest cross-section towards the ER with a 1.2-fold larger area and L_2_ was the distance to the similarly defined cross-section towards the ER. The measurements were done using a custom-made script.

### Stereology-based approach to look for ER–NE junctions in FIB-SEM datasets

We developed a stereology-based approach to search for ER–NE junctions in FIB-SEM datasets. We downloaded the datasets of a mouse pancreatic islet cell treated with high glucose (jrc_mus-pancreas-1; voxel size (nm) of 4.0 x 4.0 x 3.4 in x, y, z), a HeLa cell (jrc_hela-2; voxel size (nm) of 4.0 x 4.0 x 5.2 in x, y, z), and a human macrophage (jrc_macrophage-2; voxel size (nm) of 4.0 x 4.0 x 3.4 in x, y, z) from OpenOrganelle (Heinrich et al., 2021; Xu et al., 2021). Segmentations of the entire nucleus were required to uniformly distribute regions to search for junctions on the nuclear surface. The segmentations of the nuclei of the HeLa and the macrophage cells were available on OpenOrganelle. For the mouse pancreatic islet cell, we segmented the nucleus automatically doing a pixel classification in Ilastik (Stuart et al., 2019; version 1.3). The automated segmentations were refined using filters and other options for binary images in FIJI so that the surface of the automatically-segmented nuclei would coincide with the NE on the raw data. For each nuclei, 52–56 points were selected at the surface of the automatically-segmented nuclei using a custom-written script. The first point was selected randomly, and the rest of points were distributed uniformly at the NE in a radial way from the randomly-positioned first point. Around each point, spherical regions with a radium of 800–1000 nm were screened manually to identify potential ER–NE junctions. For the mouse pancreatic islet cell and human macrophage, we measured the length of ER–NE junctions in all the 52 and 56 regions that were selected. For the HeLa cell, we quantified the junction length in 25 out of 52 regions. To compare ER–NE with ER–ER junctions, three-way junctions were searched in the ER at random regions of cytoplasm.

### Quantification of the width of ER–NE and ER–ER junctions in FIB-SEM datasets and in budding yeast tomograms

The widths were quantified in IMOD. For constricted junctions, the width corresponded to the length of a 2-point line traced across the neck of the junction at the sagittal plane. For non-constricted junctions, the width was measured with a 2-point line across the middle of the triangulated base of the junction at the sagittal plane. The widths of ER– NE junctions identified in budding yeast tomograms were quantified in the same way as the junctions in the FIB-SEM datasets.

### Sample size determination and statistical analysis

For electron tomography, we analysed 3 cells in late anaphase, 2 cells in early telophase, and 9 cells in interphase from one experiment. We screened the NE and the cytoplasm by tomography until we found approximately 15 ER–NE and ER–ER junctions. The exact value of the analyzed surface area and the number of junctions that we found are described in main text. The statistical analysis of junction morphology was carried out after all the data were acquired. For the FIB-SEM analysis of three cells, ER–NE junctions were inspected in 52–56 uniformly-distributed regions of interest on the NE. We picked up all the potential ER–NE junctions and did not perform any data exclusion. For each cell, we found 23–34 ER–NE junctions, and we looked for a similar number of ER–ER junctions at the cytoplasm of the same cell. Statistical analyses were performed only after all the data were obtained. Statistical analysis methods, sample sizes and p-values for each experiment are indicated in figure legends.

## ACKNOWLEDGEMENTS

This project was supported by laboratory startup funding from the Medical University of Vienna to S.O. and by the Vienna Science and Technology Fund (WWTF; project LS19-001) to S.O.. H.B.T. received a DOC Fellowship of the Austrian Academy of Sciences (no. 25951) and a Max Perutz PhD Fellowship (University of Vienna and the Medical University of Vienna). We acknowledge the electron microscopy facilities at the Vienna BioCenter (Thomas Heuser, Marlene Brandstetter, Nicole Drexler, Sonja Jacob, and Harald Kotisch) and EMBL Heidelberg (especially Martin Schorb and Paolo Ronchi) for technical support and discussion. In addition, we thank Paul Wurzinger and Robert Kirmse (LEICA Microsystems) for technical support. Yeast samples were kindly provided by Anete Romanauska (Alwin Köhler’s lab). We acknowledge the members of the labs of Shotaro Otsuka (especially Clara-Anna Wagner, Tamara Völkerer, Julia Scholz, Kaike Ren, Pallavi Deolal, Pauline Vinet, and Nikoleta Kavaja), Alwin Köhler, Elif Karagöz, Roland Foisner, Peter Fuchs, David Haselbach, and Life Science Editors for feedback and discussions. Eija Jokitalo also provided us with valuable references.

## AUTHOR CONTRIBUTIONS

H.B.T. and S.O. conceived the project with input from K.I.. H.B.T. performed all the sample preparation for electron microscopy, image acquisition and analyses. S.O assisted with image acquisition. H.B.T. and K.I. established a computational image analysis pipeline to quantify ultrastructures in 3D. G.M.S. supported electron microscopy image analysis. S.O. supervised the work. H.B.T., K.I. and S.O. wrote the paper. All authors contributed to the analysis and interpretation of data.

## Notes

### Competing Interest Statement

The authors have declared no competing interest.

### Summary of Updates

Results strengthened by adding more examples of ER-NE and ER-ER junctions (Figs. 2 and 3 updated, Supplemental Figs. 1 and 4 added). New analyses performed for the junction abundance and length, and the NE width (new Fig. 4 and Supplemental Figs. 2, 3 added). Previous Fig. 4 updated (new Fig.5 and Supplemental Fig.5). Author affiliation, Introduction, Results, Discussion, Methods, and References updated.

