## Supplemental Figures for "The endoplasmic reticulum connects to the nucleus by constricted junctions that mature after open mitosis in mammalian cells"

### SUPPLEMENTARY FIGURE 1

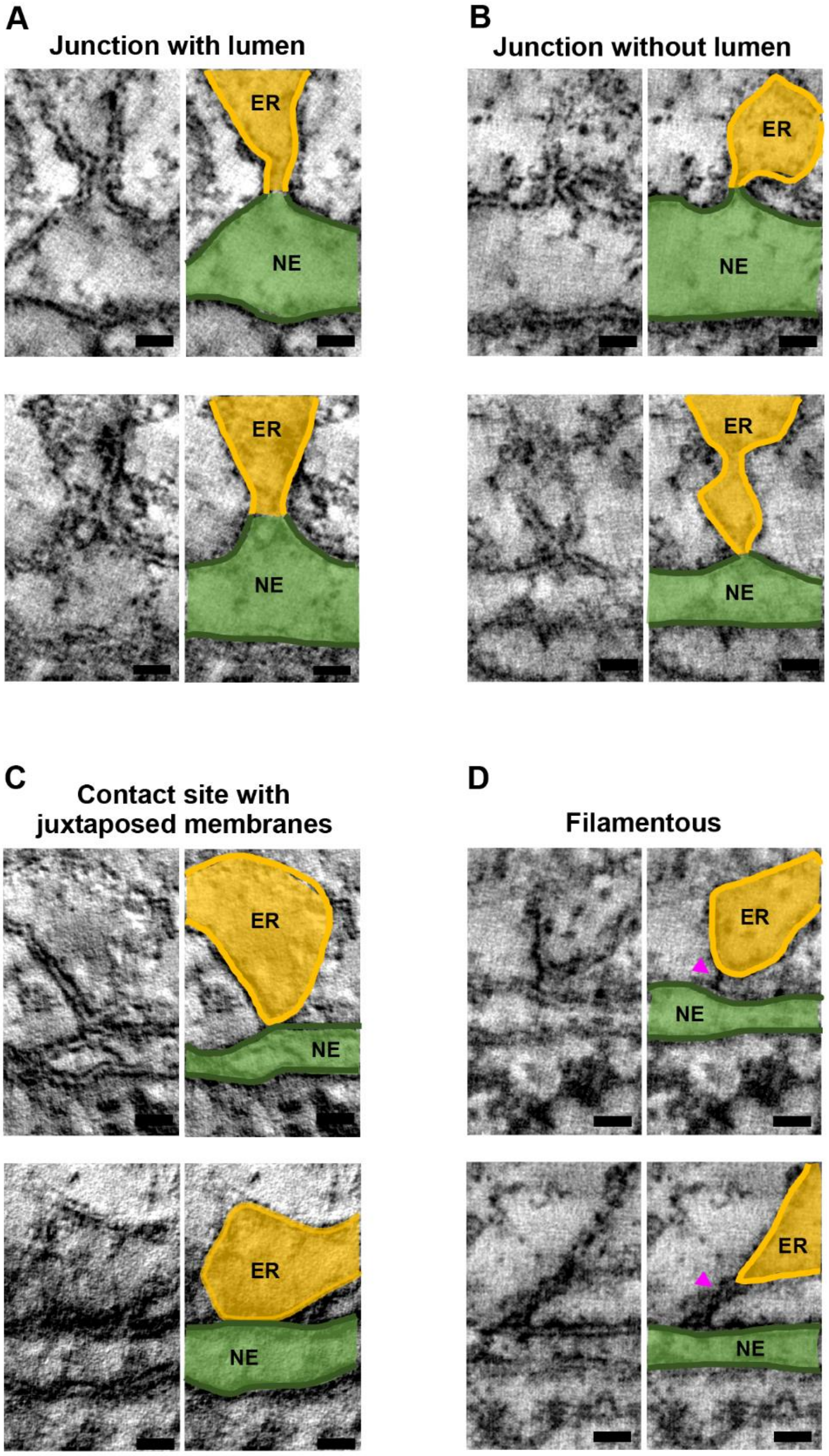

**Supplementary Figure 1. Additional examples of ER–NE junctions and different types of ER–NE contact sites in interphase. A,B** Electron tomographic slices at the sagittal plane of ER–NE junctions with (**A**) and without (**B**) lumen. For each junction, the left image shows raw EM data; the right one, the EM data on which the ER and the NE are coloured in orange or green, respectively. **C,** Electron tomographic slices at the sagittal plane of ER–NE "contact sites" where the ER membrane is juxtaposed to the ONM, but their membranes are not fused. **D,** Electron tomographic slices at the sagittal plane of "contact sites" where the ER is connected to the ONM via filamentous-like densities. The filaments are indicated with magenta arrowheads. Scale bars: 20 nm.

### SUPPLEMENTARY FIGURE 2

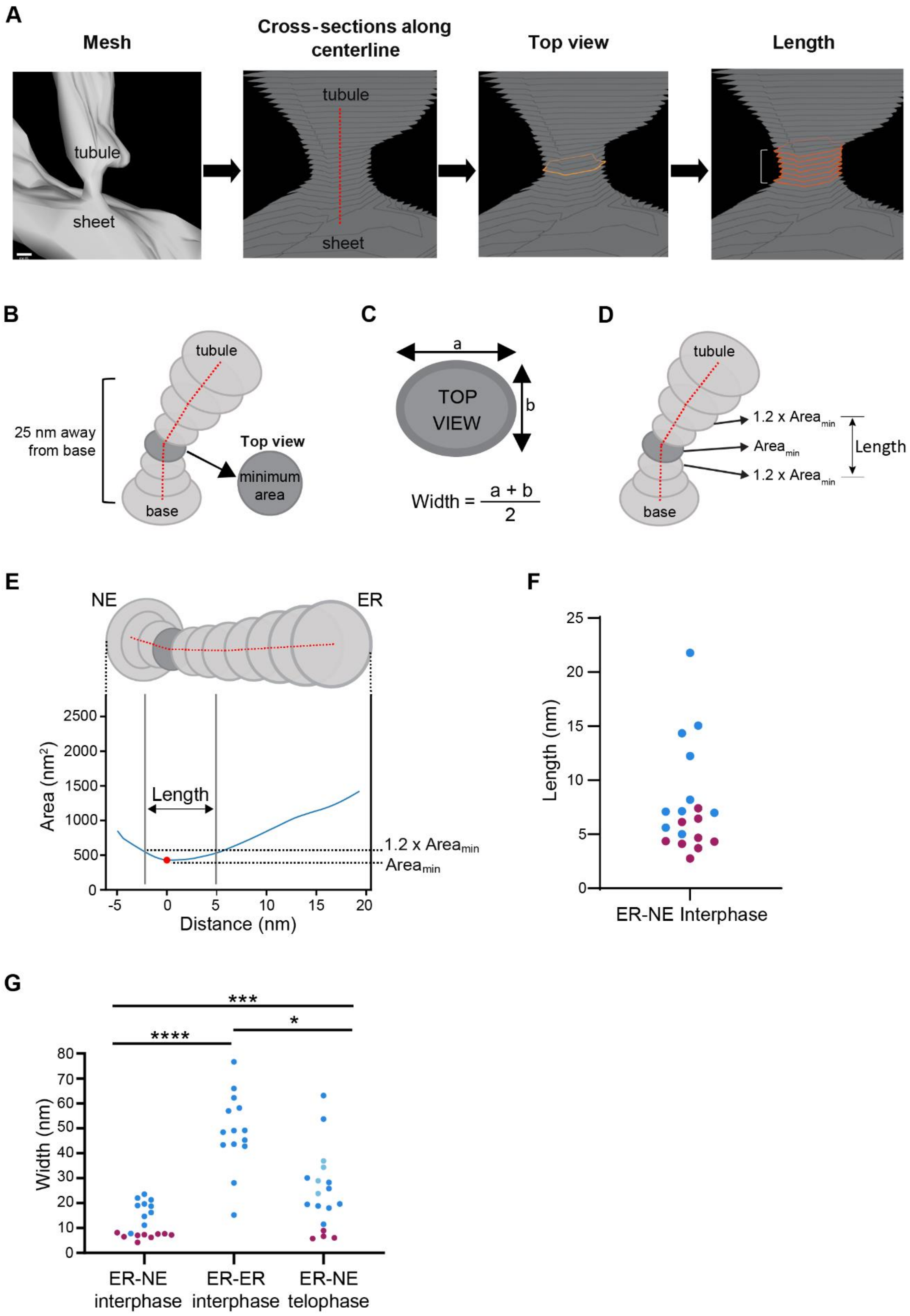

**Supplementary figure 2. 3D-ultrastructural analysis of junctions. A,** To quantify the 3D ultrastructure of the junctions, cross-sections were obtained at regularly-spaced intervals along a centreline of the 3D meshes. The cross-sections allowed identifying the junction top view and length. **B,** The cross-section with the minimum surface area was selected for making a top-view profile. **C, D,** Schemes representing how to determine the width (**C**) and the length (**D**) of junctions. **E,** A plot showing the cross-sectional area along the junction axis from the NE to the ER. The curve is centred at the cross-section with the minimum surface area. The distance between the two cross-sections with a surface area 1.2 times larger than the minimum one was defined as the length of the junctions. **F,** Length of ER–NE junctions in interphase (n = 19). The plots for the junctions with and without lumen are colour-coded in blue and magenta, respectively. **G,** Combined plots of the width of ER–NE and ER tubule-sheet junctions in interphase, as well as width of ER–NE junctions in telophase. n = 19, 14, and 18 for ER–NE interphase, ER–ER interphase, and ER–NE telophase, respectively. *p-value < 0.02, ***p-value < 0.001, ****p-value < 0.0001; two-tailed Mann–Whitney test.

#### SUPPLEMENTARY FIGURE 3

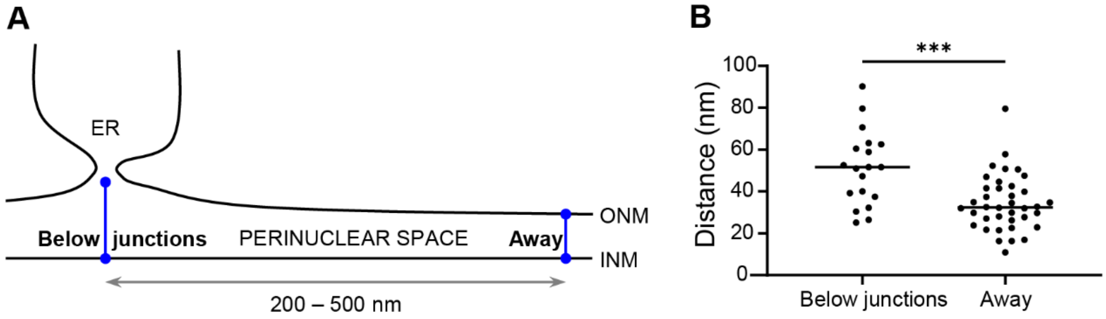


**Supplementary figure 3. Local NE dilation below ER–NE junctions. A,** A scheme depicting that the width of the perinuclear space was measured below ER–NE junctions and 200–500 nm away. **B,** Quantification of the NE width below ER–NE junctions (with and without lumen, n = 19) and 200–500 nm away from each junction (n = 38). ***p-value < 0.001; two-tailed unpaired t-test.

### SUPPLEMENTARY FIGURE 4

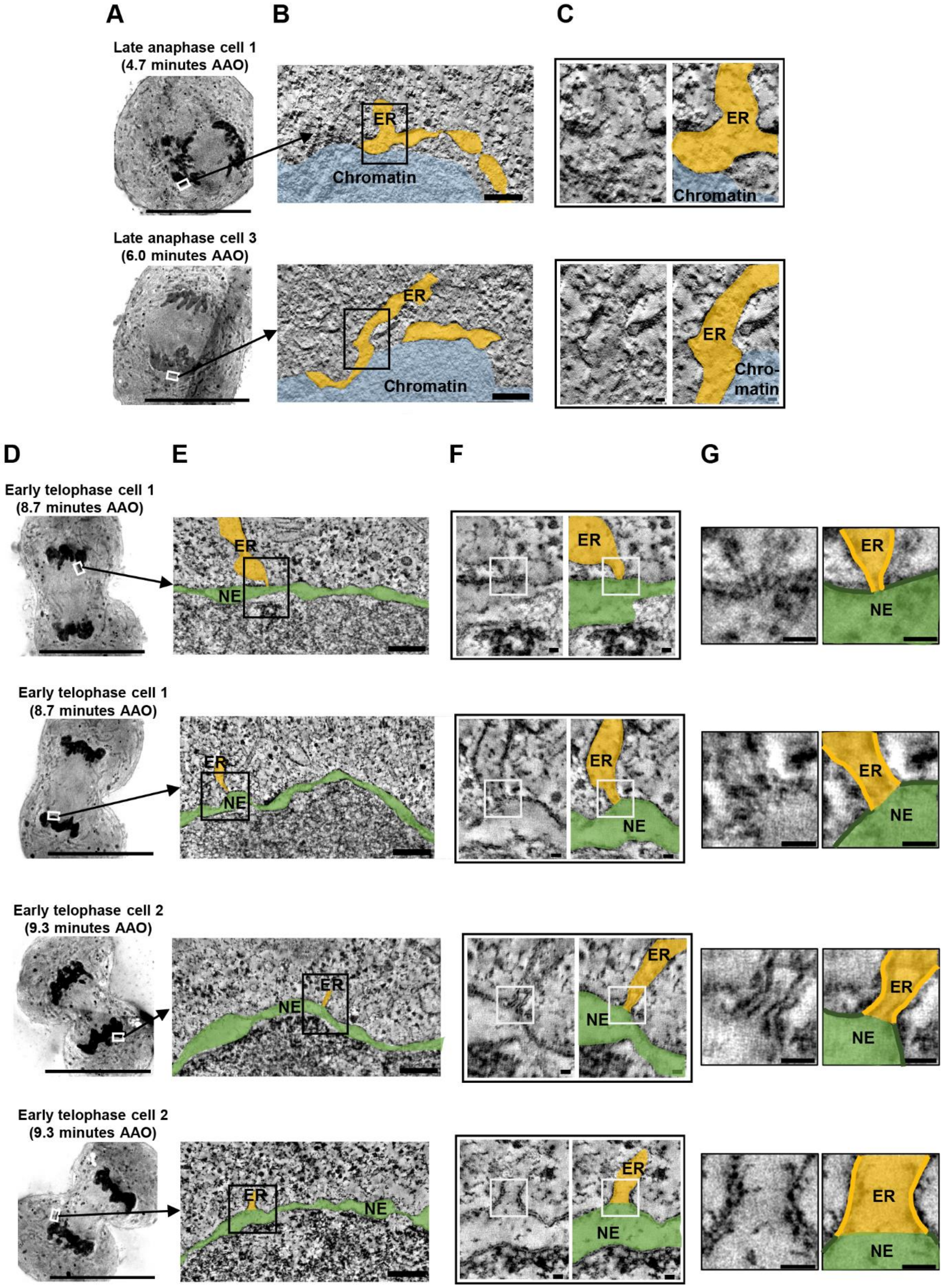

**Supplementary figure 4. Additional examples of junctions in late anaphase and in early telophase. A–C,** Additional example images of ER junctions contacting the chromatin in late anaphase cells. AAO: After Anaphase Onset. Images are displayed in the same way as in Figure 3A–C. **D–G,** Additional example images of ER–NE junctions in early telophase. Images are displayed in the same way as in Figure 3D–G.

#### SUPPLEMENTARY FIGURE 5

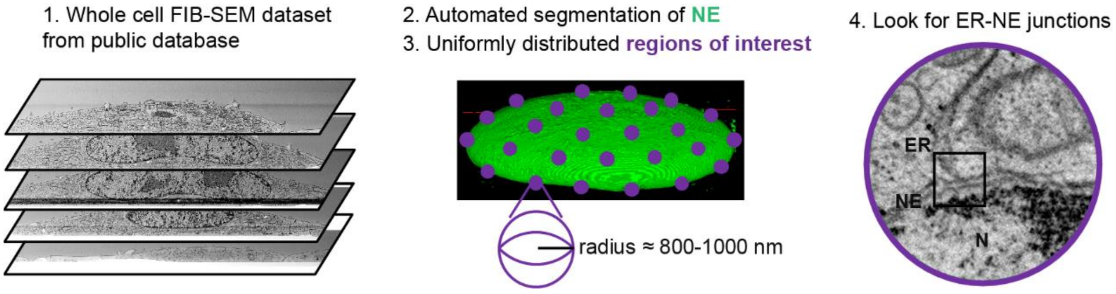


**Supplementary figure 5. Stereology-based approach to find ER–NE junctions in FIB-SEM datasets of entire cells.** The datasets were downloaded from OpenOrganelle (Heinrich et al., 2021; Xu et al., 2021). On the surface of automatically-segmented nuclei, the regions of interest were uniformly sampled with a radius of 800–1000 nm. In these regions, potential ER–NE junctions were searched manually.

#### SUPPLEMENTARY MOVIE 1

**Supplementary movie 1. Visualization of ER–NE junction in 3D with tomography.** The movie shows a junction connecting the endoplasmic reticulum to the nuclear envelope in a raw EM tomogram, as well as the 3D mesh of the junction. Note the constricted junction neck at the interface between the ER and the NE. Scale bar: 20 nm.
